# Regenerating aggregates of hydra display unique cytoskeletal organisation that is absent in a regeneration-deficient strain

**DOI:** 10.1101/2022.01.11.466547

**Authors:** Hemalatha Bhagavan, Sujana Prabhu, Niraimathi Govindasamy, Yashoda Ghanekar

**Author notes:** **Corresponding author**, Dr Yashoda Ghanekar, DeepSeeq Bioinformatics, Ananthapura Road,Yelahanka, Bangalore 560064.

## Abstract

Hydra has the unique ability to regenerate from aggregates of dissociated single cells that lack positional information. We compared two strains of hydra, a strain of hydra that was capable of regenerating from aggregates and a strain of hydra that was deficient in this type of regeneration. We observed unique actin cytoskeletal arrangements that were present in the regenerates of regeneration-competent strain but not in the regeneration-deficient strain. Concomitantly, the regeneration-deficient strain failed to organise the extracellular cytoskeleton of laminin and collagen between ectodermal and endodermal epithelial cells. These interesting preliminary observations highlight the importance of the cytoskeletal organisation in regeneration of hydra and suggest that regeneration from the aggregates of dissociated cells through *de novo* patterning requires correct structural organisation of cytoskeletal elements.

## Introduction

Axis formation or patterning followed by morphogenesis is an essential part of embryogenesis. Most organisms lose this capacity once the embryonic development is complete. Hydra is a unique system where the ability to undergo patterning and morphogenesis is retained in the adult state. A cluster of dissociated single cells from hydra can undergo self-organisation and regenerate into a functional adult^1^. Thus, starting from an aggregate of dissociated cells which does not have any positional information, the cells reorganise themselves, initiate axis formation and eventually regenerate into a functional adult polyp. Due to this unique ability, hydra is an excellent system to study regeneration and *de novo* patterning.

While performing regenerations assays from the aggregates of dissociated cells, we observed a consistent lack of regeneration in the aggregates of the strain *H. vulgaris* Ind-Pune^2,3^. Comparative studies of this regeneration-deficient strain and hydra strains capable of regeneration revealed unique cytoskeletal arrangements that were absent in the regeneration-deficient strain. This preliminary study highlights the importance of correct structural organisation of cytoskeletal elements for regeneration from aggregates of dissociated cells of hydra. Further studies will help understand the importance of the unique cytoskeletal arrangements in regeneration of hydra from aggregates of dissociated cells.

## Materials and Methods

### Hydra Culture

*H. magnipapillata* and *H. vulgaris* Ind-Pune were cultured in hydra medium (0.1 mM KCl, 1 mM CaCl_2_, 1 mM NaCl, 0.1mM MgSO_4_, 1 mM Tris-Cl, pH 8.0) as described earlier at 18°C with day-night cycle of 12 hours^4^. Hydra polyps were fed on alternate days with freshly hatched *Artemia* naupli but were starved for one day before regeneration assays and for 2 days for the rest of the experiments.

### Dissociation and regeneration of whole animal

Regeneration by aggregation of dissociated cells was performed as described earlier^1^. Briefly, head and foot regions were removed, and body columns were collected in dissociation medium (3.6 mM KCl, 6 mM CaCl_2_, 1.2 mM MgSO_4_, 6 mM Sodium Citrate, 6 mM Sodium Pyruvate, 6 mM Glucose, 12.5mM HEPES, pH 7.1, Streptomycin 100 μg/mL and Kanamycin 50 μg/mL). Body columns were washed three times with dissociation medium and then incubated in fresh dissociation medium for 30 minutes on ice. Cells were dissociated by mechanical disruption using a glass pipette. Dissociated cells were centrifuged at 200g for 10 minutes at 4°C and pellets were resuspended in dissociation medium. Cells were counted and 4×10^5^ cells were used to form one aggregate by centrifugation at 400g, 10 minutes at 4°C. Aggregates were incubated at 18°C. Dissociation medium was gradually diluted using hydra medium and was completely replaced with hydra culture medium after 24 hours. The culture medium was changed every day during regeneration.

### Generation of chimeric aggregates of *H. vulgaris* Ind-Pune and *H. magnipapillata*

To generate chimeric aggregates with ectoderm of one species and endoderm of another species, ectodermal and endodermal epithelial cells were isolated from *H. magnipapillata* as described earlier^5^ with some modifications; using solution A (equal volumes of hydra medium, dissociation medium, and 3% procaine, pH adjusted to 4.5) and solution B (equal volumes of hydra medium, dissociation medium, and 3% procaine, pH adjusted to 2.5). For the isolation of the ectoderm, the body columns were treated with solution A for five minutes, followed by solution B for 1-2 mins at 4°C and then were transferred to dissociation medium. Under these conditions, the ectoderm contracted into a ring-like structure that was separated from the rest of the body column. For the isolation of endoderm, the body columns were treated with solution A for five minutes followed by incubation in solution B for five minutes at 4°C and then transferred to dissociation medium. Ectoderm disintegrated and separated from endoderm under these conditions and endoderm was isolated for further experiments. Isolated ectoderm and endoderm were either used as intact layers or were dissociated mechanically to generate single cell suspensions. Endoderm and ectoderm of *H. vulgaris* Ind-Pune were isolated by a similar procedure as above, but with 1% procaine solution instead of 3%.

### Immunostaining

For all the staining procedures, aggregates were fixed with 4% paraformaldehyde (PFA) for 30 minutes at room temperature followed by permeabilization for 30 minutes in 0.1% Triton-X. To stain actin filaments, aggregates were incubated with phalloidin (1:50) conjugated with FITC (Molecular Probes, Invitrogen) for 1h at room temperature. For immunostaining, aggregates were blocked with 1mg/ml BSA for 1h at room temperature and then incubated with mouse anti-laminin antibody (1:100) or mouse anti-collagen 1 (1:100) antibody at 4°C for 12-14 hours^6^. Unbound antibodies were washed thrice with PBS and incubated with anti-mouse Alexa 647 (1:200) for 1h at room temperature. All the aggregates were washed thrice with PBS and stained with 1 μg/mL 4’,6-diamidino-2-phenylindole (DAPI) for 10 minutes at room temperature. Aggregates were mounted in mowiol with 2.5% 1,4-diazabicyclo[2.2.2]octane (DABCO). Images were captured at 20X for laminin and collagen 1 immunostaining and at 60X for actin cytoskeleton using the Zeiss LSM 550 confocal microscope or using Zeiss LSM 700 confocal microscope equipped with ZEN imaging Software (Zeiss, Germany).

### Mass spectroscopy

Aggregates of *H. vulgaris* Ind-Pune and *H. magnipapillata* strains were used for total protein isolation. Aggregates were lysed by incubating in lysis buffer containing 10 mM Tris-Cl, pH 7.5, 1 mM sodium chloride, 1 % Triton X-100, and 10 μg/ml Protease Inhibitor Cocktail (Roche). The aggregates were then sonicated for 2 minutes, followed by centrifugation at 10000rpm for 30 minutes at 4°C to collect cellular proteins. The supernatant was collected and frozen at −80°C till use.

Protein lysates were electrophoresed on 12% polyacrylamide gel. After electrophoresis, the gel was stained using 0.5% Coomassie stain for two hours at room temperature. Gel was destained for one hour using 50% methanol, 40% water and 10% acetic acid and then imaged using the GE ImageQuant system. A single band above 250KDa which was completely absent in the aggregate protein lysate of *H.vulgaris* Ind-Pune but present in *H. magnipapillata* was given for mass spectrometric analysis.

## Results

### Regeneration defect in the *H. vulgaris* Ind-Pune

Hydra body columns can be dissociated mechanically to generate single cell suspension, and clusters of these single cells regenerate into a functional adult^1^. These aggregates of dissociated single cells from *H. magnipapillata* can regenerate into functional polyps as reported earlier and shown in figure 1, left panel. During regeneration, the ectodermal and endodermal epithelial cells in the aggregate reorganise to form two distinct layers with a smooth outer surface within 4-6 hours after aggregation. Hollow cavities develop in aggregates around 12-16 hours, which merge by 24 hours. Tentacle buds appear by 48 hour and rudimentary hydranths with hypostome and tentacles develop in 72-96 hours after aggregation. We call this strain regeneration-competent (RC) strain. While performing similar regeneration assays in the strain *H. vulgaris* Ind-Pune^2, 3^, we found a consistent lack of regeneration from the aggregates. In the aggregates of *H. vulgaris* Ind-Pune, ectodermal and endodermal epithelial cells initiated reorganisation in 4-6 hours. A clear outer layer with a dark core, indicating separation of ectodermal and endodermal layers was visible in the aggregates by 16h (Figure 1, 16h time point of the regeneration-deficient strain). However, the aggregates failed to develop beyond this stage (Figure 1, right panel) and disintegrated by 48 hours. We call *H. vulgaris* Ind-Pune strain regeneration-deficient or RD strain.

**Figure 1.**
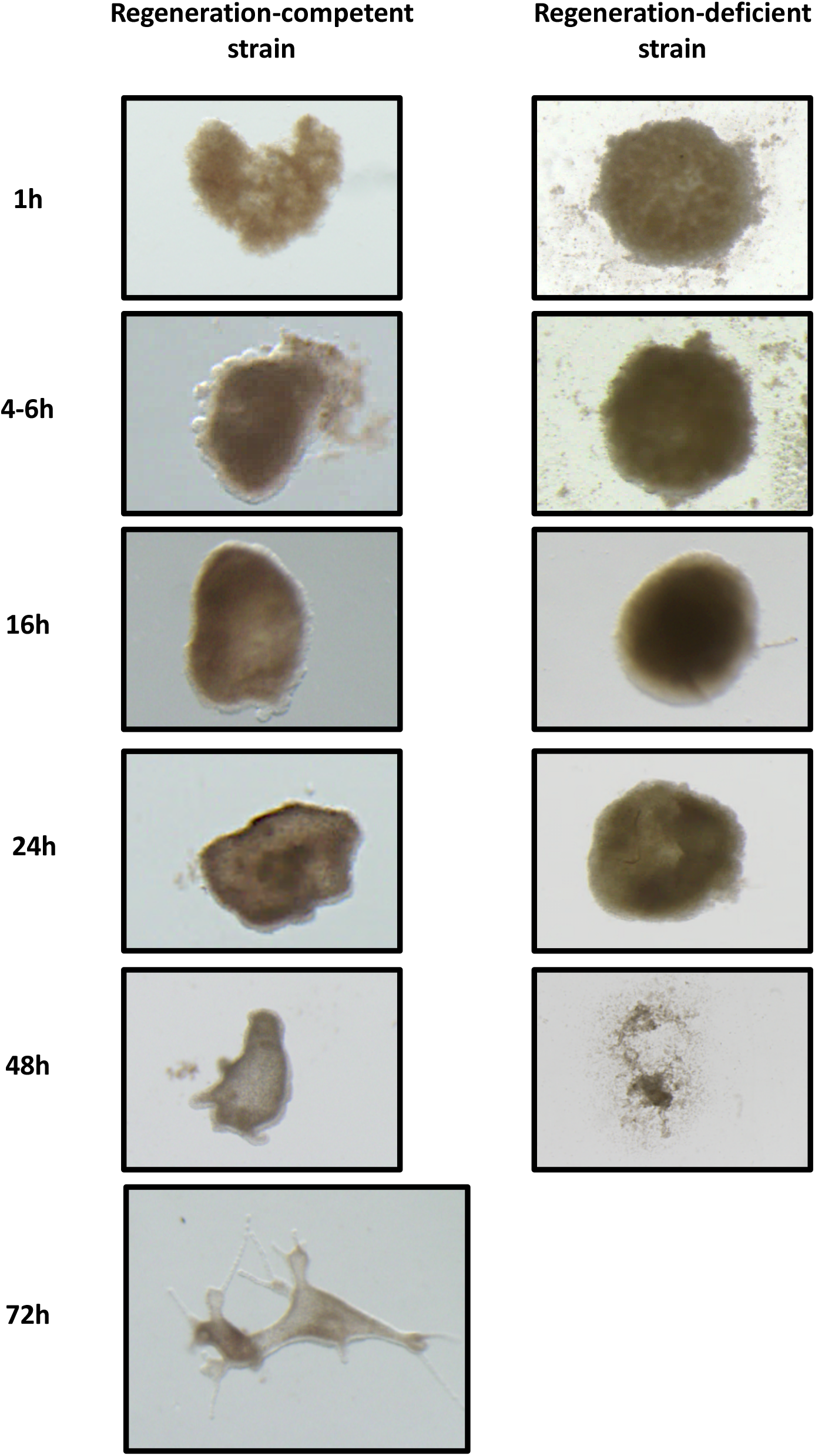
Regeneration of Dissociated Cells of Hydra: The left panel shows different stages of regeneration of aggregates of dissociated single cells from *Hydra magnipapillata,* a strain of hydra that is capable of this type of regeneration. The right panel shows the fate of aggregates of dissociated cells from *H. vulgaris* Ind-Pune or the RD strain, which is not capable of undergoing this type of regeneration.

### Regeneration defect lies in the endodermal layer of *H. vulgaris* Ind-Pune

Regeneration defects observed previously in hydra head regeneration have been attributed to specific defects in different cell types^7–9^. Both ectodermal and endodermal epithelial cells, but not interstitial cells, are involved in the establishment of the epithelial bilayer from the unstructured mass of single cells, followed by establishment of axis in the aggregate^5, 10^. It is possible that either or both epithelial layers could be deficient in the processes specifically involved in this type of regeneration from aggregates but not in regeneration after mid-gastric bisection or in normal physiology. To test this hypothesis, we generated chimeric aggregates consisting of ectodermal and endodermal epithelial cells from RC and RD strains. Ectodermal and endodermal epithelia can be isolated and dissociated to obtain single cell suspensions of ectodermal epithelial cells and endodermal epithelial cells^5^. In these assays, the epithelial cells undergo reorganisation to generate an inner endodermal epithelial layer and an outer ectodermal epithelial layer followed by regeneration.

We generated chimeric aggregates using dissociated cells from isolated epithelial layers^5^. Equal numbers of ectodermal and endodermal epithelial cells were mixed to generate chimeric aggregates using ectoderm from one species and endoderm of another species. Ectodermal and endodermal epithelia of the same species were combined as control. In aggregates with both ectodermal and endodermal epithelial cells from RC strain, the epithelial cells underwent rearrangement to form a bilayer, followed by regeneration into functional hydranths, in a manner similar to regeneration of aggregates of whole body columns. The aggregates with ectodermal and endodermal epithelial cells from RD failed to develop, as expected from the previous experiments. While the epithelial cells reorganised to generate a bilayer with ectoderm covering the endoderm, the aggregate failed to regenerate further (Table 1).

**Table 1.** Regeneration from chimeric aggregates. Ectodermal and endodermal layers of regeneration-competent and regeneration-deficient strains were isolated as described under Methods. The layers were dissociated to generate single cell suspension. Chimeric aggregates were generated by mixing equal numbers of ectodermal cells and endodermal cells of the two strains and regeneration was monitored. Aggregates with endodermal cells from regeneration-deficient strain failed to develop. On the other hand, aggregates with ectodermal cells from regeneration-deficient aggregates were able to undergo regeneration.

| Source of Ectoderm | Source of Endoderm | Regenerated aggregates/Total seeded aggregates |
| --- | --- | --- |
| Regeneration-competent | Regeneration-competent | 9/9 |
| Regeneration-deficient | Regeneration-competent | 8/9 |
| Regeneration-competent | Regeneration-deficient | 0/9 |
| Regeneration-deficient | Regeneration-deficient | 0/9 |

Interestingly, the aggregates with endodermal epithelial cells of RD, when combined with ectodermal epithelial cells of RC did not regenerate. In contrast, the aggregates that were generated using dissociated ectodermal epithelial cells of RD in combination with endodermal epithelial cells of RC were able to regenerate (Table 1). These studies suggested that the lack of regeneration was due to a defect in the endoderm of the RD strain. This lack of regeneration was not due to incompatibility of cells belonging to two different species, as the aggregates of ectodermal epithelial cells of RD and endodermal epithelial cells of RC were able to regenerate.

### Actin and extracellular matrix in regenerating aggregates

The reorganisation of ectodermal and endodermal epithelial cells in aggregates involves migration of ectodermal and endodermal epithelial cells to generate a bilayer with ectodermal epithelial cells forming the outer layer and endodermal epithelial forming the inner layer. This is followed by secretion of mesoglea or the extracellular matrix between the two epithelial layers and establishment of cell-cell contacts. The bilayer formation was not affected in the aggregates of the RD strain, and the ectodermal/endodermal epithelial cells reorganised similar to those in the RC strain (Figure 1, aggregates after 16h). We therefore ruled out the possibility of a defect in the migration of epithelial cells. This observation also rules out the possibility of deficiencies or irregularities in the homotypic or heterotypic fusions that take place in regenerating aggregates^11^.

Mesoglea or the extracellular matrix in hydra is an elastic, flexible structure present between the two epithelial layers. It is composed of basal lamina-like structure in contact with each epithelial layer and it is composed of laminin and collagen type IV. Between laminin and collagen type IV are fibrillar collagens like collagen type I and proteoglycans. Newly formed aggregates of dissociated cells are devoid of extracellular matrix. Extracellular matrix can be detected between the two epithelial layers about 17 hours after aggregation and regeneration does not take place if biogenesis of extracellular matrix is perturbed^12^. Actin is another structural element important for maintaining cell morphology, polarity, migration and cell-cell adhesion. To investigate if the lack of regeneration seen in the RD stain was due to perturbation in dynamics of these structural elements involved in morphogenesis, we investigated the expression and localization of actin and mesoglea components in the regenerating aggregates of RC and RD strains.

Phalloidin staining was performed in the aggregates of RC and RD strains to investigate organisation of filamentous actin. By 24 hours, a large amount of filamentous actin could be seen in both types of aggregates. Actin filaments in the apical sections displayed a distinct actin belt-like structure (Figure 2A, top panel) in RC strain but not in RD stain (Figure 2B). Such actin belts in ectodermal epithelial cells have been reported earlier^13^. Interestingly, a closer look at the confocal sections with nuclei ~8μm below the sections with polymerised actin showed that these actin belt-like structures actually encompassed 3-4 nuclei rather than a single nucleus. Figure 2A, bottom panel shows maximum intensity projection through 8 μm from the surface of regenerating aggregates. As shown in this panel, 3-4 nuclei were surrounded polymerised actin. Since the confocal sections with nuclei were only ~8μm lower than the sections with polymerised actin, with the imaging performed from the surface of the aggregates, these nuclei belong to the ectodermal epithelial cells. These observations suggest that regenerating aggregates display a unique arrangement of actin belt-like structures that encompass 3-4 ectodermal epithelial cells rather than a single epithelial cell. While the RD strain expressed large amount of polymerised actin, it was largely disorganised with some polymerized actin observed around a single nucleus in some places, but it did not have any ordered organisation or a belt-like structure similar to regeneration-competent strain (Figure 2B).

**Figure 2.**
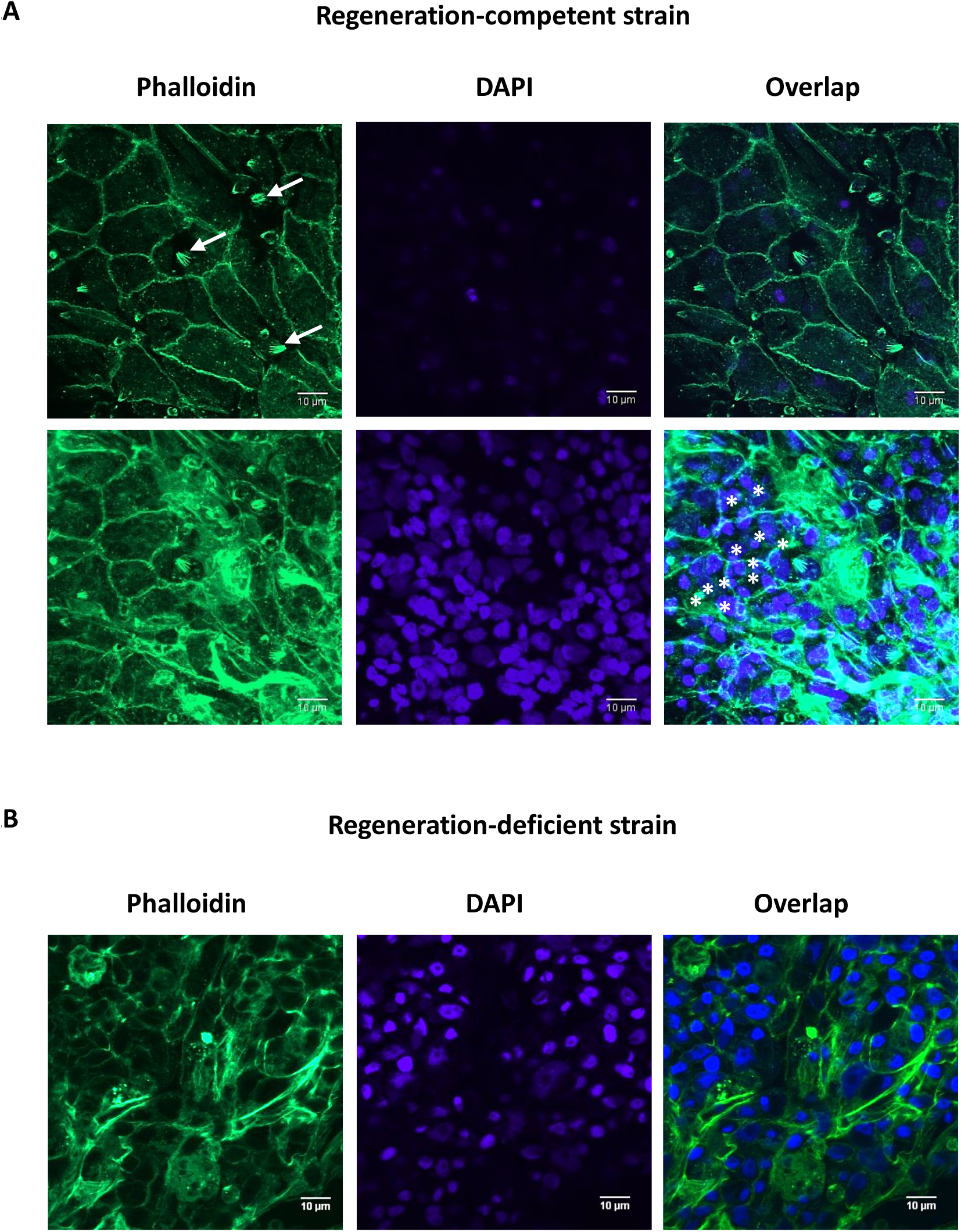
Actin Organisation in Regenerating Aggregates: (A) The figure shows phalloidin staining of regeneration-competent hydra aggregates at 24h. Belt-like structures of polymerised actin are seen in the top panel, along with punctate structures (highlighted by arrows). The bottom panel shows maximum intensity projection through 8 μm from the surface of regenerating aggregates and shows 3-4 nuclei surrounded by actin belt-like structures. The asterisks in the bottom panel show nuclei surrounded by an actin belt-like structure. (B) The figure shows phalloidin staining of aggregates from regeneration-deficient strain of hydra. There are no organised actin belt-like structures and few punctate structures.

We also observed punctate foci of filamentous actin, reminiscent of focal adhesion in the RC strain. Numerous such punctate structures were present in RC stain (Figure 2, top panel). These punctate structures were also seen in RD, but their size and numbers were substantially lower as compared to the RC strain (Figure 2B).

Laminin, one of the main components of extracellular matrix is synthesized by endodermal epithelial cells while collagen I is synthesized by ectodermal epithelial cells^14, 15^. In the RC strain, laminin localisation at the basal lamina could be observed at 24 hours as reported and was present between the two epithelial layers (Figure 3, top panel). Expression of laminin was also observed in the endodermal epithelium of the RD strain, however, laminin was not localised between the two epithelial layers (Figure 3, bottom panel). The expression and deposition of collagen between the two epithelial layers was also observed in RC strain. At 24 hours, collagen I expression and its deposition between the two epithelial cells was initiated (Supplementary figure 1, top panel) and this was more prominent at 36h (Supplementary figure 1, middle panel). In the RD strain, collagen I deposition between the two epithelial layers was not observed in at 24 hours (Supplementary figure 1, bottom panel). Collagen staining was not possible in this strain at a later time point like 36h, since these aggregates started disintegrating by this time.

**Figure 3.**
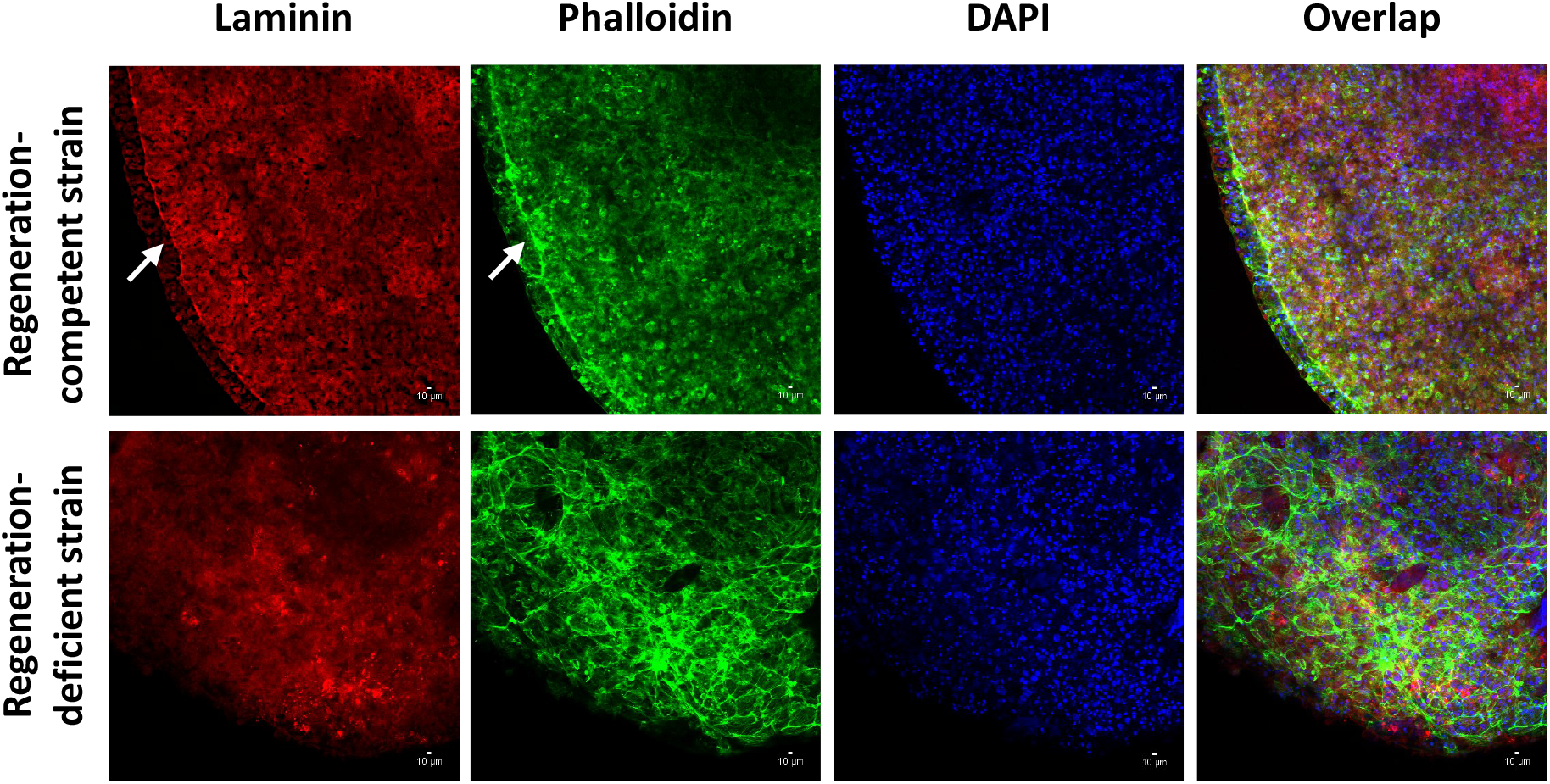
Laminin localisation in regenerating aggregates: The figure shows laminin staining of aggregates of regeneration-competent strain (top panel) and regeneration-deficient strain (bottom panel) at 24h after aggregation. While laminin is expressed by both strains, localization of laminin between ectodermal and endodermal epithelial cells is seen only in regeneration-competent strain, as highlighted by arrows. Laminin also colocalizes with actin in regeneration-competent strain.

### The RD strain aggregates do not express structural proteins

We also performed mass spectrometry in order to investigate if there are any differences in the expression of proteins between the two strains. Proteins were extracted from aggregates of RC and RD strains 24 hours after aggregation. Upon electrophoresis, we observed that a group of high molecular weight proteins that were expressed in RC, but were absent in RD (Supplementary figure 2). Using mass spectrometry, four proteins that were present in RC strain but not in RD strain were identified. These proteins include a-spectrin, ß-Spectrin, dynein Heavy Chain and microtubule actin cross-linking factor. There were however no differences in the expression levels of the transcripts of these proteins as seen by quantitative PCR (data not shown), suggesting that the differences in expression levels are due to post-transcriptional regulation.

## Discussion

Our studies on the comparison of a regeneration-deficient and regeneration-competent strains of hydra show that correct structural organisation of actin cytoskeleton, laminin and collagen is essential for *de novo* patterning followed by regeneration in hydra. Patterning does not occur in the absence of correct organisation of these structural elements, even though they are abundantly expressed. The failure of the regeneration-deficient strain to regenerate beyond the formation of epithelial bilayer is likely due to the absence of the correct organisation of cytoskeletal scaffolds of actin and the glue of extracellular matrix that keeps these cells together in the bilayer.

Importantly, the study revealed an interesting structure of actin cytoskeleton where polymerised actin was found to encircle a group of 3-4 cells. This is similar to the actin belts found in the apical epithelium. However, instead of encircling one ectodermal epithelial cell, this belt-like structure was found to encompass 3-4 ectodermal epithelial cells. The regeneration-deficient strain lacked these structures. To our knowledge, such actin belts around several cells have not been reported earlier in hydra or any other system. Actin belts have been observed in hydra by Aufshnaiter et al, but these were reported to be present around a single cell^13^. However, these images do not have corresponding nuclear staining and it is not clear if the actin belt shown in Aufshnaiter *et al* encompasses more than one cell.

It is also possible that this arrangement of actin polymers is specific to regenerating aggregates. Such an organisation of epithelial cells might form a functional unit that could be important in regeneration. Clusters of 3-4 cells could be essential for eventual formation of signalling centres such as the head organiser in hydra development. For example, such a unit of 3-4 cells might allow concentration of a signalling molecule or transcription factor to build up inside the cluster, which can lead to development of a gradient for *de novo* axis formation.

This study also revealed another structure that has not been reported in hydra so far. The regenerating hydra aggregates showed the presence of punctate actin structures resembling focal adhesions. These structures are present in large numbers in the strain capable of regeneration. In the RD strain, they are smaller and fewer in number. The role of these structures is not clear. Concomitantly, the RD strain also lacked proteins required for crosslinking of cytoskeletal elements such as Spectrin Alpha, Spectrin Beta, Dynein Heavy Chain and Microtubule Actin cross-linking factor as seen in mass spectrometry. It is possible that the cytoskeletal components that are not expressed in aggregates of RD strain are required for the formation of these structures. It is possible that the missing elements and structures are essential for initiating the organisation of actin cytoskeleton that is required to hold together the two epithelial layers and the mesoglea, since this strain of hydra otherwise appears normal and has been used extensively for studies involving regeneration of foot and head regions^3^.

Observations from this study indicate the importance of cytoskeletal organisation for *de novo* axis formation in hydra. However, these studies require further investigations to confirm that the nuclei surrounded by an actin belt-like structures belong to different cells using plasma membrane markers and to rule out presence of several nuclei within belt-like structures is due to phagocytosis. Electron microscopic analysis can further help validate the observations made in this study such as differences in the organization of ectodermal and endodermal epithelial cells as well as cytoskeletal elements in the two strains. Comparative studies investigating actin cytoskeletal structures in polyps and regenerating aggregates would also help in elucidating the role of cytoskeletal structures in hydra regeneration.

It is possible that the *H. vulgaris* Ind-Pune strain lost this ability for *de novo* generation of actin cytoskeleton during evolution. Since this ability is not essential for reproduction by budding or regeneration (except in laboratories investigating the phenomenon of regeneration from aggregates of dissociated single cells), it did not affect its survival.

## List of Abbreviations

RC: regeneration-competent
RD: regeneration-deficient

## Acknowledgements

Authors thank Surendra Ghaskadbi for providing the strain *H. vulgaris* Ind-Pune and Xiaoming Zhang, University of Kansas, USA for providing laminin and collagen antibodies. Authors also thank the Central Imaging and Flow Facility and Mass Spectrometry Facility at the Bangalore Biocluster for confocal imaging and mass spectrometry respectively. Authors would like to thank the members of YG laboratory at the Institute of Stem Cell Biology and Regenerative Medicine for discussions and inputs.

## Author contributions

YG conceived, designed and supervised the project. All experiments pertaining to hydra aggregations were performed by HB and SP. Confocal imaging was performed by NG. YG, HB, SP and GN analysed the data. YG wrote the manuscript and HB made the figures.

