## Supplemental Figures for "Regenerating aggregates of hydra display unique cytoskeletal organisation that is absent in a regeneration-deficient strain"

### Supplementary Figure 1

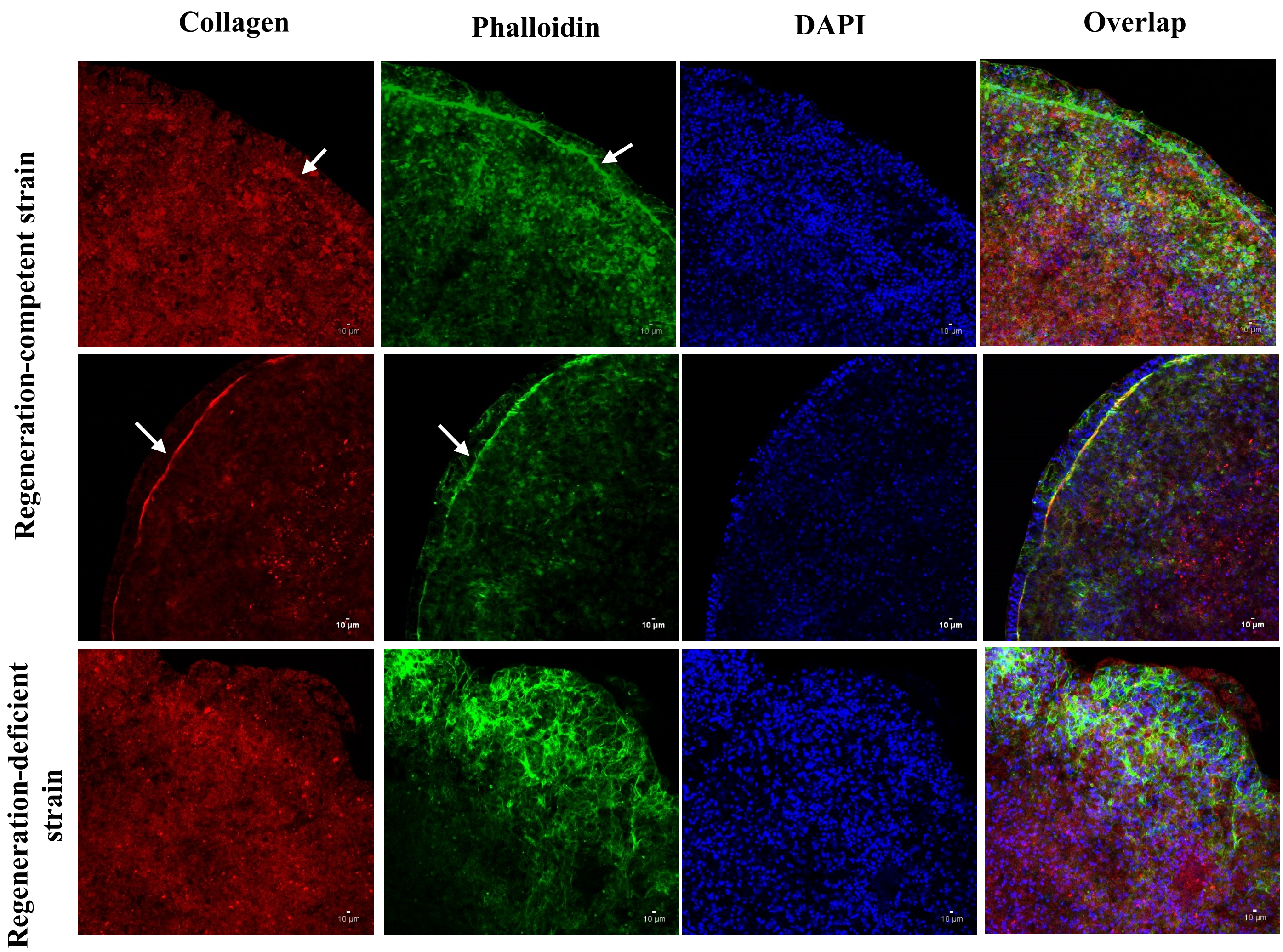


**Supplementary Figure 1**

**Collagen localisation in regenerating aggregates.** The figure shows collagen staining of aggregates of regeneration-competent strain (top panel) and regeneration-deficient strain (bottom panel) at 24h after aggregation. Collagen is expressed by both strains at 24h. Collagen staining of regeneration-competent strain at 36h is shown in the middle panel. Deposition of collagen between ectodermal and endodermal epithelial cells is initiated at 24h (top panel, shown by arrow), and is more prominent at 36h (middle panel, shown by arrow). The regeneration-deficient strain does not show any deposition of collagen between ectodermal and endodermal epithelial cells.

**Supplementary Figure 2**

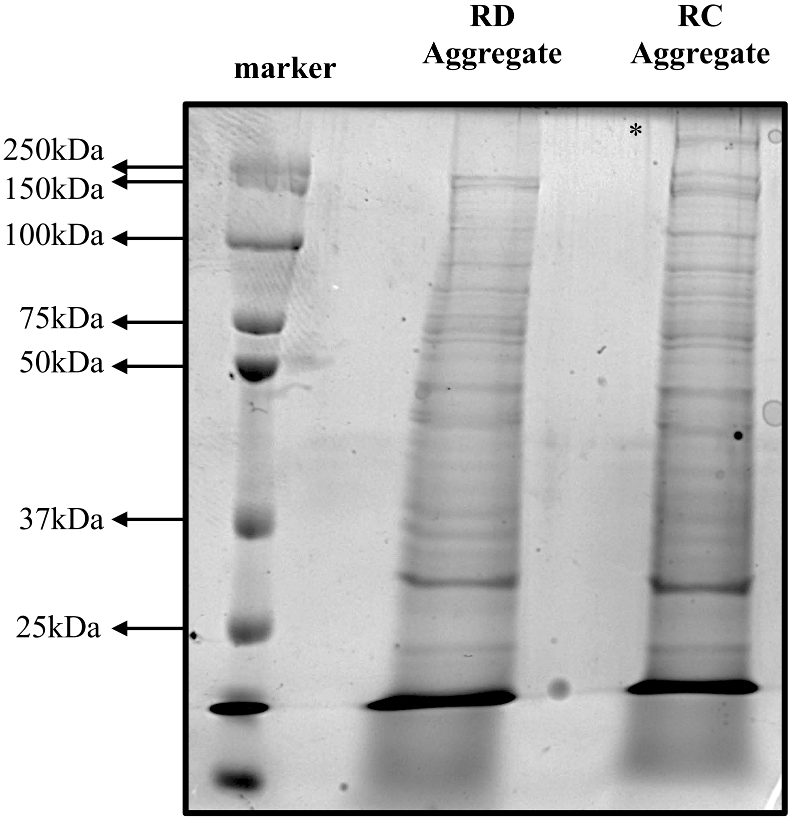


**Supplementary Figure 2**
**Protein expression in regeneration-competent and regeneration-deficient aggregates.** The figure shows Coomassie-stained gels of protein lysates of regeneration-competent and regeneration-deficient aggregates at 24h. A prominent high molecular weight protein band (>250kDa, marked by asterisk) which was present in regeneration-competent strain but not in regeneration-deficient strain was given for Mass Spectrometry.
